# Structural insights into G protein activation by D1 dopamine receptor

**DOI:** 10.1101/2022.01.18.476830

**Authors:** Xiao Teng, Sijia Chen, Qing Wang, Zhao Chen, Xiaoying Wang, Niu Huang, Sanduo Zheng

## Abstract

G protein-coupled receptors (GPCRs) comprise the largest family of membrane receptors and are the most important drug targets. An agonist-bound GPCR engages heterotrimeric G proteins and triggers the exchange of GDP with GTP to promote G proteins activation. A complete understanding of the molecular mechanisms of G proteins activation has been hindered by a lack of structural information of GPCR-G protein complex in nucleotide-bound states. Here, we present the cryoelectron microscopy (cryo-EM) structures of D1 dopamine receptor (D1R)-G_s_ in the nucleotide-free state, the GDP-bound state and the GTP-bound state with endogenous ligand dopamine. These structures reveal important conformational changes accounting for the release of GDP and the GTP-dependent dissociation of Gα from Gβγ subunits. Combining mutagenesis functional studies, we also identified an important sequence motif in D1R that determines its G protein selectivity. Taken together, these results shed light into the molecular basis of G protein selectivity and the entire molecular signaling events of GPCR-mediated G protein activation.

---

G protein-coupled receptors (GPCRs) mediate numerous physiological functions by responding to a wide range of stimuli including light, odors, hormones and neurotransmitters (*1*). Agonist binding to a GPCR induces its conformational changes which subsequently lead to the engagement of guanosine diphosphate (GDP)-bound Gαβγ heterotrimer. Structural rearrangement of Gα when bound to GPCR results in the exchange of GDP for guanosine triphosphate (GTP) and the dissociation of heterotrimer. Gα are divided into three major subfamilies: adenylyl cyclase stimulatory G protein (Gα_s_), adenylyl cyclase inhibitory G protein (Gα_i/o_) and Gα_q/11_ on the basis of distinct downstream signaling pathways. Most GPCRs couple primarily to one type of Gα. Understanding the molecular mechanisms of G protein activation and selectivity has been the subject of intensive research. The first crystal structure of the β2-adrenergic receptor (β2AR)-G_s_ complex in the nucleotide-free state revealed outward movement of TM5 and TM6 in β2AR when coupling to G protein compared to the inactive β2AR, which creates a large cytosolic pocket of β2AR (*2*). The C-terminal helix (α5) of Gα_s_ displaced towards the receptor and inserted into the cytosolic pocket of the β2AR. The conformational changes of the GPCR-G protein interface allosterically induce structural rearrangement of the nucleotide-binding pocket, leading to the separation of the α-helical domain (AHD) of the Gα subunit from the Ras-like domain (Ras) and the subsequent release of GDP. In complement to structural studies, hydrogen/deuterium exchange mass spectrometry (HDX-MS) (*3, 4*), double electron-electron resonance spectroscopy (DEER) (*5*) and molecular dynamics (MD) studies (*6*) have shown that both the AHD and Ras domain separation and the conformational change of the nucleotide-binding pocket caused by GPCR-G protein interaction are necessary to promote the GDP release.

Since the report of the first crystal structure of β2AR-Gs complex, an increasing number of structures of GPCRs-G proteins complex were obtained by single particle cryo-electron microscopy (cryo-EM) (*7, 8*). These are attributable to the use of scaffold proteins (*2, 9, 10*) to stabilize the GPCR-G protein complex and modified thermostable G proteins (mini-G) (*11*), and the technical breakthroughs in cryo-EM (*12*). However, all of these complex structures solved so far are in the nucleotide-free state, which only provide a snapshot of a stable intermediate state. The GPCR-G protein coupling events are obviously highly dynamic and comprise a series of intermediate states. A recent crystal structure of β2AR in complex with a C-terminal peptide of Gα_s_ revealed a different configuration from the β2AR-G protein complex, providing additional insights into the molecular basis of G protein selectivity (*13*). Clearly, it is important to obtain intermediate states of GPCR-G protein complex including GDP and GTP-bound state at atomic level in order to fully understand the molecular mechanisms of G protein selectivity and G protein activation. However, instability of the GPCR-G protein complexes in the nucleotide-bound state makes them intractable to structural studies.

Dopamine exerts a variety of physiological functions through five distinct G protein-coupled dopamine receptors subtypes (D1R to D5R), including locomotor activity and reward (*14–16*). Dysfunction of the dopaminergic system has been linked to Parkinson’s disease and psychiatric diseases. DRs are classified as two subfamilies: the D1-like (D1R and D5R) and the D2-like (D2R, D3R, and D4R). Although DRs share high sequence similarity in the transmembrane region involved in G protein binding, D1-like receptors couple to Gs, while D2-like receptors couple to G_i/o_ **(Fig. 1A)**. Recently published cryo-EM structures of D1R-G_s_ and D2R-Gi_/o_ with various ligands provided structural insight into ligand recognition and G protein selectivity (*17–21*). In this study, to better understand the molecular basis of G protein selectivity and activation, we sought to determine the cryo-EM structures of the D1R-Gs complex in both nucleotide-free and nucleotide-bound states.

**Fig. 1.**
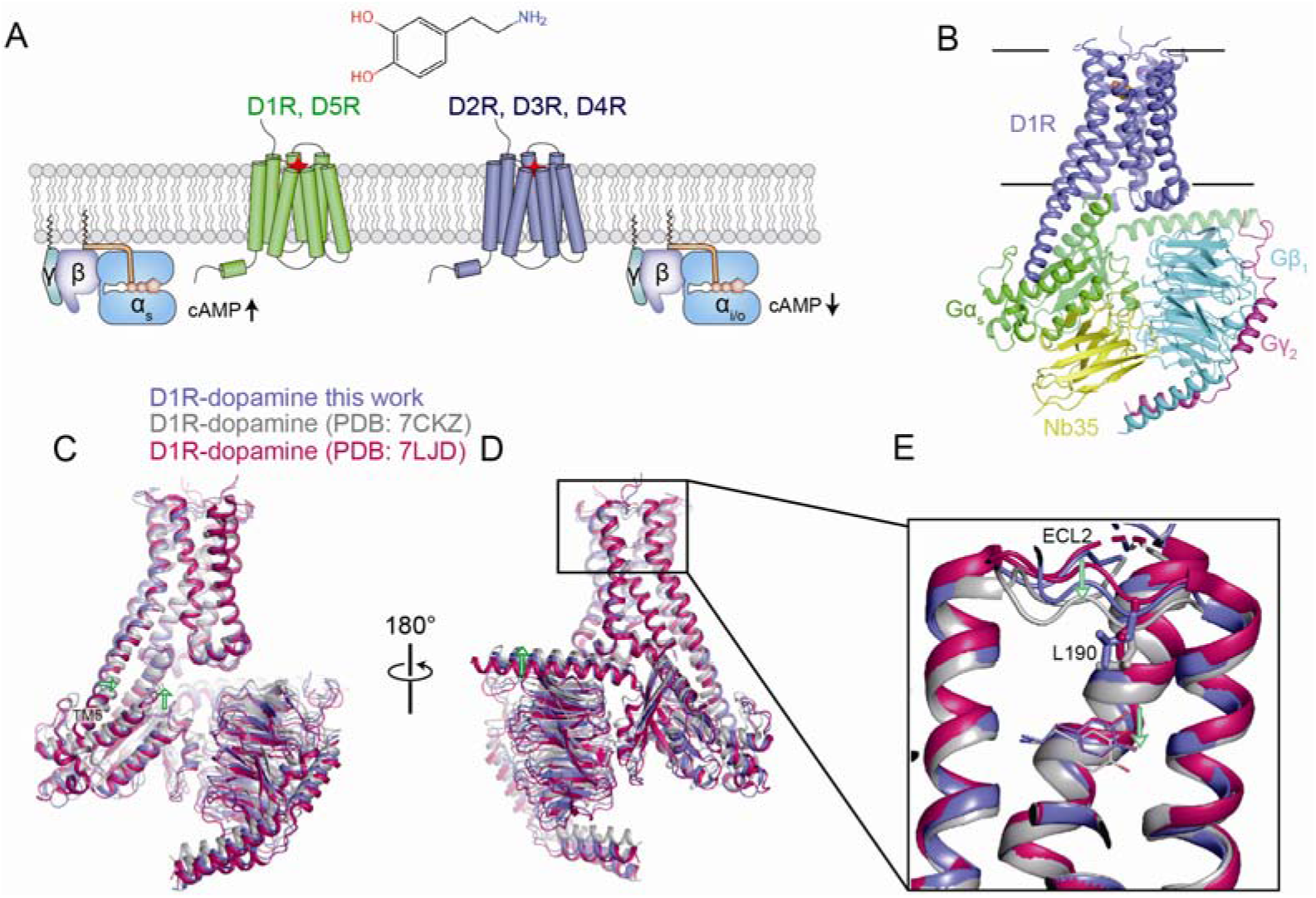
Structure of the dopamine-bound D1R-G protein complex in the nucleotide-free state. **(A)** G protein coupling selectivity among dopamine receptors. **(B)** Overall architecture of dopamine-bound D1R-miniGs-Nb35 complex. D1R, Gα_s_, Gβ_1_, Gγ_2_ and Nb35 are colored in blue, green, cyan, magenta and yellow respectively. **(C and D)** Structural superposition of the dopamine-bound D1R-G protein structure in this study and dopamine-bound structures of the same complex in previous studies in two opposite views. Conformational changes were shown with green arrows. **(E)** Close-up views of the dopamine binding pocket. L190 at ECL2 involved in hydrophobic interaction with dopamine was shown as stick.

## Structures of dopamine-bound D1R-mini-G_s_ complex

To enhance the stability of D1R-G_s_ complex and simplify the purification process, we created a fusion protein (D1R-mini-Gα_s_) where the C-terminus of the wild-type human D1R is fused to the N-terminus of mini-Gα_s_ (*11*) which is an engineered thermostable G_s_ without the AHD domain. We expressed D1R-mini-Gα_s_ in Expi293 cell by transiently transfection and purified it by antibody affinity chromatography. To assemble the D1R-mini-G protein complex, the purified D1R-miniGα_s_ was mixed with the excess Nb35 that has been used to stabilize the GPCR-G protein complex and human Gβ_1_γ_2_ subunits and further purified to homogeneity by size-exclusion chromatography **(fig. S1A)**. Structures of the dopamine-bound D1R-mini-G_s_ complex in the nucleotide-free state, GDP-bound state and the GTP state were determined at nominal resolutions from 3.1 to 4.2 Å **(fig. S, 1 to 6 and table S1)**. Small molecules including dopamine and GDP except GTP can be unambiguously modeled owing to the excellent quality of EM density map. Due to the high stability of the D1R-miniGs fusion protein complex and no orientation preference, we were able to obtain structures at atomic resolution with around 600 movies. Moreover, D1R can form a stable complex with G protein without Nb35 **(fig. S3)**.

The overall arrangement of the D1R-miniG_s_-Nb35 complex is largely similar to the previously determined GPCR-G_s_ protein complex **(Fig. 1B)**. The high stability of the D1R-Gs complex may be attributed to the more extensive interaction interface between D1R and Gα than that between β1AR and Gα, including 2.5 helical turns of TM5 extension **(fig. S2A)**. When compared to the β1AR-G_s_ complex, the entire Gαβγ heterotrimer in the D1R-G protein complex is rotated clockwise relative to the receptor **(fig. S2, A and B)**. As a result, D312 at Gβ subunit is in close proximity to K339^8.52^ at helix 8 of D1R, leading to a close contact between Gβγ and D1R **(fig. 2C)**. The TM5 extension in D1R likely accounts for the distinct orientation of the receptor and G protein from the β1AR-G_s_ complex (*21*). These findings suggest that the relative orientation of the receptor and G protein is very dynamic and may vary during the GPCR-G protein coupling cycle.

**Fig. 2.**
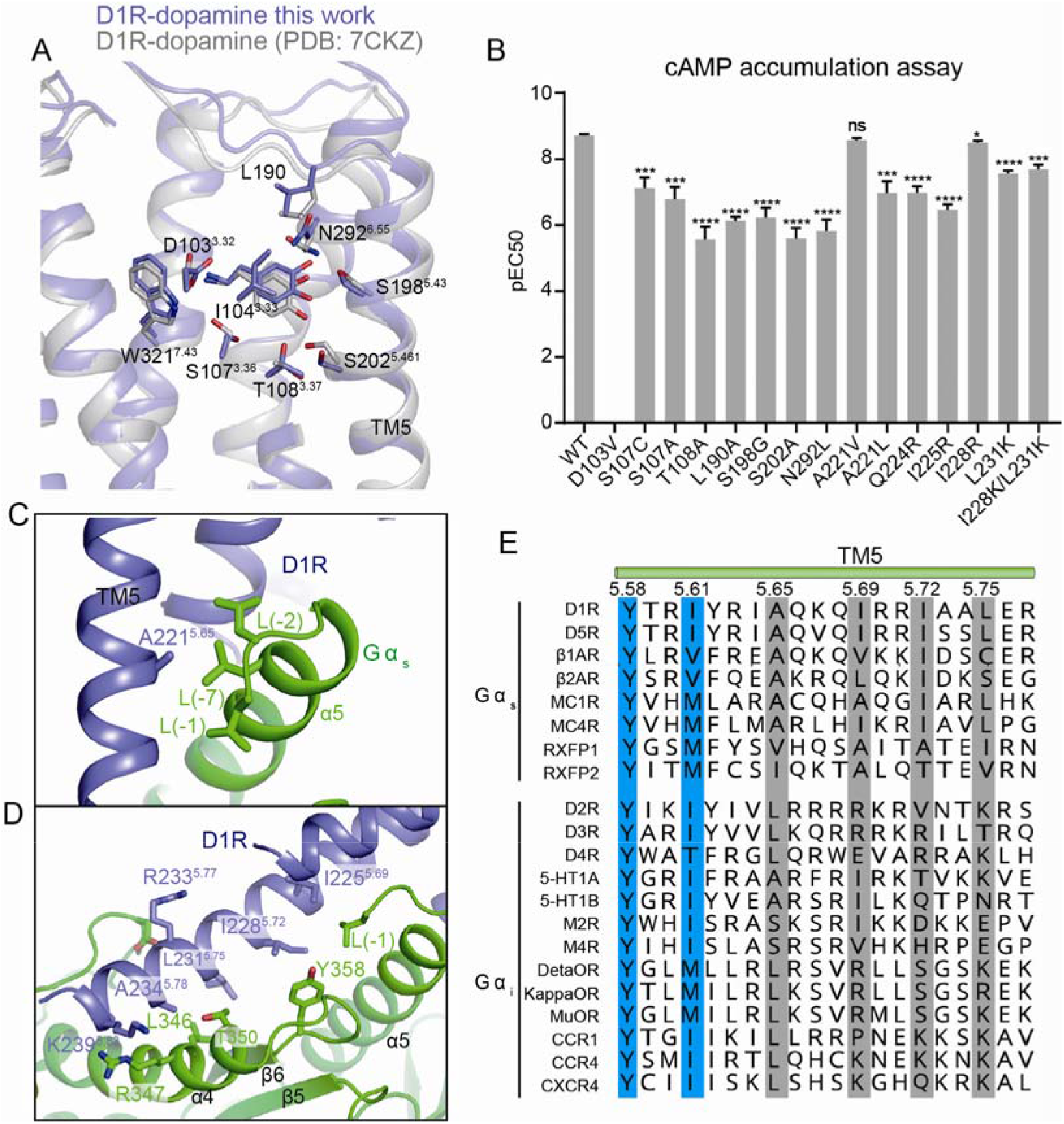
Molecular determinants of the G protein selectivity by dopamine receptors. **(A)** Comparison of the binding pose of dopamine between our structure and the previously determined structure (PDBID: 7CKZ). **(B)** cAMP accumulation assay of D1R and D1R mutants activated by dopamine. **(C)** A221^5.65^ of the receptor engages hydrophobic interactions with L388, L393 and L394 at the α5 of Gα. **(D)** Detailed interactions between the TM5 extension and Gα. **(E)** Sequence alignment of the C-terminal part of TM5 from several G_s_-coupled receptors and G_i/o_-coupled receptors.

## Plasticity of the ligand binding site

Interestingly, when comparing our structure with two recently published structures of dopamine bound D1R-Gs complex (*18, 19*), we found that the binding pose of dopamine varied among these structures **(Fig. 1, C to E**). While the binding modes of amine groups of dopamine which make salt bridge interaction with D103^3.32^ (Superscript corresponding to the Ballesteros-Weinstein numbering system) are almost identical, the catechol ring moves downwards. The downward movement of the catechol ring in the binding pocket is accompanied by an upward shift of the entire Gαβγ and the inward movement of TM5 **(Fig. 1, C and D)**. In our structure, S198^5.43^ makes strong hydrogen bonds with both hydroxyl groups of catechol, and the para hydroxyl group is distant from and engages weak hydrogen bond interactions with both S202^5.46^ and T108^3.37^ compared to the previously reported structure (PDB ID: 7CKZ) (*19*) **(Fig. 2A)**. The downward movement of the catechol ring makes the para hydroxyl group close to the S202^5.46^ and T108^3.37^ in TM5 **(Fig. 2A)**, allosterically leading to further inward movement of TM5 and upward shift of G protein **(Fig. 1, C and D)**. L190 in ECL2 moves in the same direction as dopamine, suggesting it plays an important role in dopamine binding **(Fig. 1E)**. The functionally equivalent residue of L190 in D2R is I184 which neighbors L190 and is located above dopamine when aligning two structures **(fig. S2D)**. Consistent with our structural observations, mutation of any residues involved in binding dopamine significantly reduced G protein coupling efficiency **(Fig. 2B)**. Previous studies have shown that G protein coupling to the receptor allosterically influences the conformation of the agonist binding pocket (*22*). Therefore, the conformational differences of the D1R-G protein interface among different studies that may arise from different versions of G protein used for structural studies lead to the conformational heterogeneity of the ligand binding pocket and the different binding mode of dopamine. The different binding pose of the same ligand has also been observed between two D2R-G_i_ complex structures determined in micelle and lipid environment respectively, which is also attributed to conformational differences of the interface of the receptor and G protein (*17*). Taken together, these results suggest that the conformation of the ligand binding pocket and the binding pose of ligands vary depending on the conformation of the cytoplasmic side of the receptor that may change during the receptor-G protein coupling process or through interaction with different downstream effectors.

## The importance of the C-terminal part of TM5 in determining G protein specificity

The important role of ICL2 especially the hydrophobic residue at position 34.51 in determining Gs coupling selectivity has been well studied (*19, 23, 24*). In this work, we focused on the other regions that contribute to G protein selectivity of DRs. Most of residues in TM3, TM5 and TM6 involved in interactions with Gs are conserved in D2R **(fig. S2, E and F)**. Notably different residues are located at the C-terminal part of TM5 including TM5 extension (**Fig. 2, C and D**). For example, A221^5.66^ is projected into a hydrophobic pocket formed by L(−7), L(−2) and L(−1) of α5 in Gα_s_ (−1 represents the last residue of Gα_s_) **(Fig. 2C)**. While most G_s_-coupled GPCRs prefer hydrophobic residues with smaller side chains including valine and alanine than leucine at the equivalent position of A221^5.66^, G_i_-coupled GPCRs can accommodate a variety of hydrophobic residues including leucine **(Fig. 2E)**. Substitution of A221^5.66^ to valine in D1R had little influence on the potency of dopamine, whereas substitution of leucine resulted in significantly reduced potency **(Fig. 2B)**. From a structural perspective, A221^5.66^L mutation likely leads to steric clashes with the aforementioned hydrophobic pocket of α5 in Gα_s_ due to their close distance. In addition, three hydrophobic residues including I225^5.69^, I228^5.72^ and L231^5.75^ are located at the C-terminus of TM5, and form extensive hydrophobic interactions with the Ras domain of Gα_s_. The three equivalent residues are hydrophobic residues in most G_s_-coupled GPCRs, whereas at least one of the three equivalent residues in G_i_-coupled GPCRs is a charge residue including lysine or arginine **(Fig. 2E)**. Mutations of I225^5.69^ into charge residues significantly impaired the potency of dopamine, and the effect of I228^5.72^ or L231^5.75^ mutation was modest **(Fig. 2B)**. The charge residues are particularly enriched in the C-terminus of TM5 in Gi-coupled receptors, and have been shown to be critical for Gi coupling (*25*). The important roles of A/V^5.66^ and I225^5.69^ in determining Gs selectivity were further verified using NanoBiT-based assay which can directly assess effects of these mutations on interactions between D1R and Gs **(fig. S3, G and H)**. Moreover, the coupling efficiency between D2R and Gs was dramatically enhanced when the ICL3 in D2R including the motif was substituted by that in D1R **(fig. S3I)**. Similarly, G_i_-coupled α2 adrenergic receptor acquired the ability to activate G_s_ by replacing its ICL3 with that of the β2AR (*26*). Collectively, these results indicate that the A/V^5.65^Φ^5.69^ motif (Φ represents hydrophobic residues) in TM5 is predominant in Gs-coupled receptors, and plays an important role in determining Gs selectivity.

## Structural basis for the GDP release upon G protein activation

Structures of GPCR-G protein complexes in the nucleotide-free state have shown that receptor binding to Gα_s_ allosterically induces conformational changes of the α5-β6 loop, α1 and P loop of the nucleotide binding site in Gα as well as the separation of the AHD from the Ras domain, which are critical for receptor-mediated nucleotide release (*7, 27*). However, it is yet to be determined as to the conformational steps of G protein activation and which regions are the major determinant for the initial release of GDP (*28*). To answer these questions, we sought to determine the structure of the D1R-G protein complex in the presence of GDP. The overall structure of the GDP-bound D1R complex in the present of Nb35 is similar to that of the D1R-G protein complex in the nucleotide-free state **(Fig. 3A and fig. S3, A to D)**. To rule out the possibility that Nb35 restricts the conformational change of the complex caused by GDP binding, we also determined the structure of GDP-bound D1R-G protein complex without Nb35 **(Fig. 3B and fig. S3, E to G)**. GDP were well-defined in EM densities map of GDP-bound D1R-G protein complex with or without Nb35 **(fig. S3D and S4A)**. The switch II of Gα undergoes large conformational change in the absence of Nb35, leading to a roughly 2 Å translational movement of the Gαβγ towards TM5, suggesting that Nb35 actually influence the relative orientation of the receptor and G protein by stabilizing the conformation of the switch II **(fig. S4, B and C)**. Compared to the GDP-bound Gα_s_ without receptor binding, GDP-bound Gα_s_ in D1R-G complex shares common structural changes with the D1R-G complex in the nucleotide-free state in α5 of Gα_s_, which undergoes rotational and translational movement **(Fig. 3C)**. Structural studies of the GPCR-G protein complex in the nucleotide-free state suggest that ICL2 binding to the G protein induces the conformational change of the αN-β1 hinge region, which is propagated to the P loop through β1, the conformational change of which results in GDP release (*7*). However, our structure show that the conformation of P loop and α1 involved in binding of the diphosphate of GDP almost remain in place upon receptor binding prior to GDP release, whereas V367 in the α5-β6 loop move away from GDP by about 3 Å because of the structural rearrangement of α5 when engaged by the receptor **(Fig. 3D)**. Since V367 sandwiches GDP with K293 in αG, and is also involved in interaction between AHD and Ras domain **(Fig. 3E),** V367 movement weakens both the interaction between Gα and GDP and the interaction between the AHD and the Ras domain. Previous mutagenesis studies have shown that insertion of a flexible linker including five glycine residues but not a rigid alpha-helical segment between TCAT/V motif (T/V corresponds to V367 in Gα_s_) and α5 blocks the G protein activation by GPCRs (*29*). This flexible linker absorbs the structural change of α5 induced by receptor binding and disrupts the conformational change of V367, which eventually prevents GDP release. To further support our structural observations, we performed in vitro GTP-turnover assay using the purified D1R and G_s_ heterotrimer. As expected, D1R catalyzed rapid GDP/GTP exchange on Gα subunits, compared to the G_s_ heterotrimer alone, and the GTP-turnover rate of D1R for the V367A mutant of Gα_s_ was substantially increased **(Fig. 3G)**, underscoring the important role of V367 in receptor-induced GDP release. Moreover, another noticeable feature in the structure of the GDP-bound complex is the rotational movement of α1 in Gα **(Fig. 3F)**, which possibly plays a key role in the separation of the AHD domain from the Ras domain. In the GDP-bound Gα without receptor binding, F376 of α5 engages aromatic interactions with H64 of α1, F212 of β2 and F219 of β3, and Q59 of α1 makes hydrogen bonds with T369 of α5 **(Fig. 3F)**. When engaged by F129^34.51^ in ICL2 of D1R, F376 in α5 undergoes translational and rotational movement, which disrupts its aromatic interactions with nearby residues and the hydrogen bond between Q59 and T369, leading to the translational movement of F212 and F219 and the rotational movement of H64 and Q59 in α1 (*30*) **(Fig. 3F)**. The movement of Q59 causes a steric clash with L198 in AHD, thus destabilizing the AHD-Ras domain interface. The functional importance of F^34.51^ in ICL2 was shown by a mutation to alanine that significantly reduced the GTP-turnover rate of D1R **(Fig. 3G)** and almost abrogated GDP release induced by β2AR (*3*). Besides, the slower GTP-turnover rate of the family B glucagon receptor could be attributed to the absence of strong hydrophobic interactions between the residue in ICL2 analogous to F^34.51^ in D1R and β2AR, and Gα_s_ (*31*). Furthermore, the steric effect of Q59 was supported by mutagenesis studies showing that the GTP-turnover rate of D1R in Q59L mutant of Gα_s_ but not Q59A mutant was dramatically increased. This can be explained by the fact that although both Q59A and Q59L mutants disrupt the hydrogen bond between Q59 in α1 and T369 in α5, alanine fails to mimic the steric effect of Q59 due to its smaller side chain. Moreover, T369A mutation in Gα_s_ had little effect on GTP-turnover rate of D1R **(Fig. 3G),** whereas the equivalent mutation, T329A in Gα_i_ caused a significant increase in receptor-independent GDP release (*32*). Taken together, our results indicate that receptor binding to Gs protein induces the rotational movement of Q59 in α1 that causes the separation of AHD from Ras, and the conformational change of V367 in the α5-β6 loop that weakens GDP binding, both of which are critical for G protein activation. Following GDP release prior to GTP binding, the α1 and α5-β6 loop move further towards the TM5 of the receptor, while the α5 remains in place **(fig. S4D)**. The conformational dynamics of α1 and the α5-β6 loop during G protein activation are also demonstrated by HDX-MS results showing that receptor binding induced an increase in HDX in these regions (*3*).

**Fig. 3.**
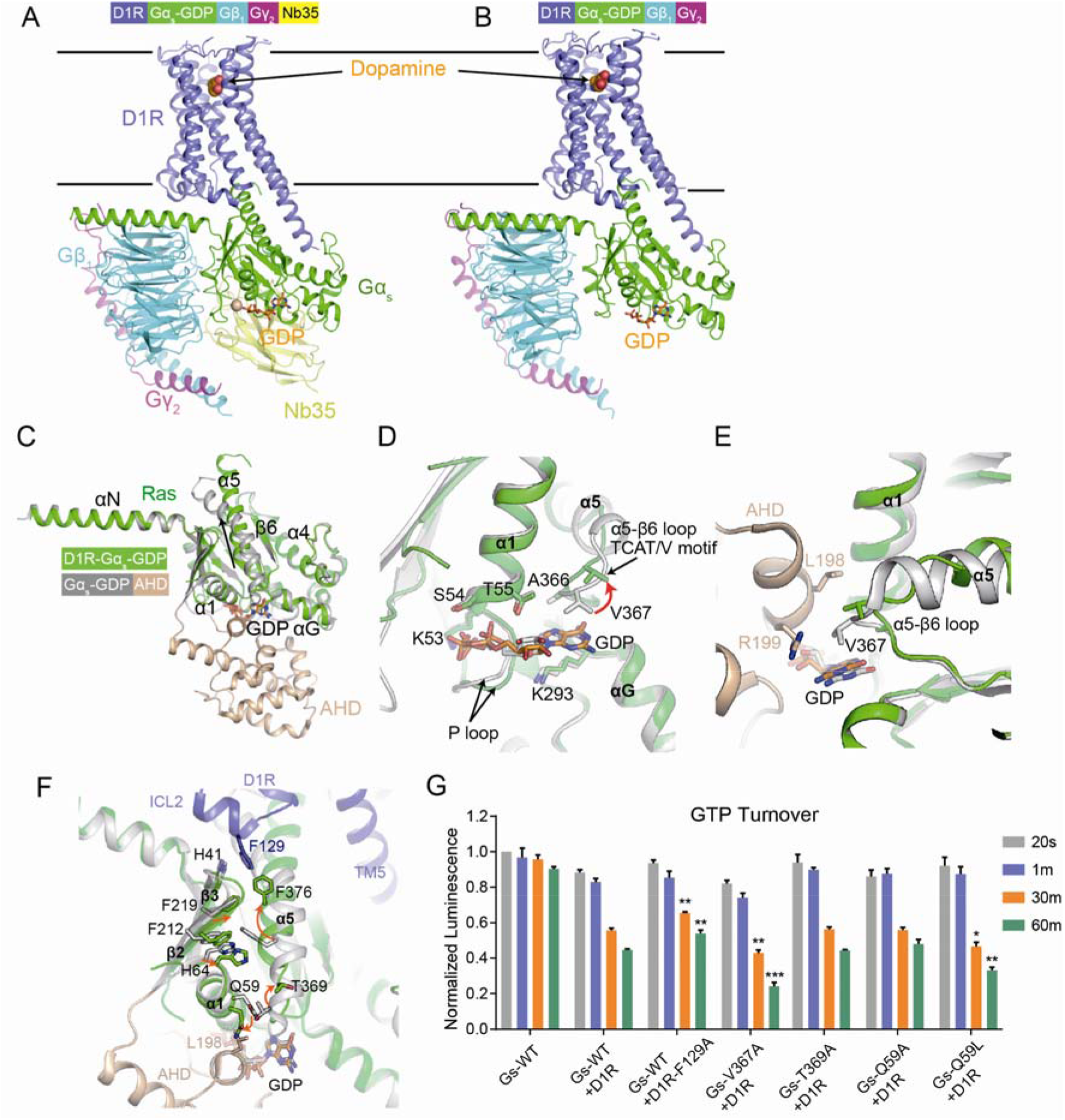
Structural changes of Gα upon receptor engagement prior to GDP release. **(A)** Structure of dopamine-bound D1R-mini-Gs-Nb35 complex in the presence of GDP. GDP was shown as sticks and colored in orange. The same color scheme as Figure 1b was used for proteins. **(B)** Structure of the dopamine-bound D1R-miniGs complex without Nb35 in the presence of GDP. **(C)** Comparison of the structures of receptor-free Gα_s_ (PDB ID: 6EG8) and D1R-bound Gα_s_ (green) in the presence of GDP. D1R and Gβγ were omitted for clarity. The Ras domain and α-helical domain (AHD) in free Gα_s_ are colored in grey and wheat, respectively. **(D)** The receptor induces the conformational change of α5 which subsequently leads to the upward movement of V367 at α5-β6 loop. **(E)** Structural change of V367 influences the interaction between AHD and Ras. **(F)** The conformational change of α5 leads to structural arrangement of α1, which disrupts the interaction between AHD and Ras domain. **(G)** GTP turnover experiments of WT Gs or mutants induced by D1R receptor. Significance is calculated by comparing the wild type and mutants at the same time point using two-tailed student’s t-test.

## Structure of GTP-bound D1R-G protein complex

Although the structure of GTP-bound Gα has provided insight into mechanisms of the GTP-dependent dissociation of Gα from Gβγ (*33*), it remains unclear how GTP triggers the dissociation of G proteins from receptors. The mini-Gα_s_ we used for structure determination includes an I372A mutation at α5 which makes the receptor-G protein complex resistant to GTP-mediated dissociation (*34*). We speculate we may capture a GTP-bound intermediate state prior to the receptor-G protein dissociation. Indeed, D1R can form a stable complex with G protein in the presence of GTP from the 2D classification **(fig. S5A)**. We were able to obtain two different structures, one with Nb35 occupied and one with Nb35 dislodged after 3D classification **(Fig. 4A and fig. S5, B to F)**. The γ-phosphate of GTP interacts with the switch II of Ras domain and leads to its structural arrangement, which subsequently expels the Nb35 **(Fig. 4, B and D)**. The conformational change of the switch II arising from GTP binding causes the movement of Gβγ by about 3.8 Å **(Fig. 4, C and D)**. In contrast, the αN-β1 hinge in Gα moves by only 1 Å, because of strong hydrophobic interactions between F129^34.51^ in ICL2 of D1R and residues in the αN-β1 hinge, the β2-β3 loop and α5, which limits the movement of the αN-β1 hinge. As a result, the imbalanced movement of Gβγ and the αN-β1 hinge in Gα disrupt the interface of αN and Gβγ, such that the αN helix of Gα in the GTP-bound D1R-G complex is tilted around 20 degrees towards the receptor compared to that in the D1R-G complex in the nucleotide-free or GDP-bound state **(Fig. 4B)**. The movement of αN results in smaller interaction interface between Gα and Gβγ in the GTP-bound D1R-G protein complex **(Fig. 4E)**. Moreover, GTP binding causes the displacement of H41 in the αN-β1 hinge and F219 in β3 away from α5, enlarging the hydrophobic pocket where F129 is inserted, and weakening interactions between Gα and D1R **(Fig. 4F)**. The movement of αN observed in our structure is consistent with results of fluorescence labeling experiments and HDX-MS showing that αN underwent large conformational change upon interaction with receptors and GTP (*3, 24, 35, 36*). However, the conformational change of αN was not captured in previous structural studies of GPCR-G protein complexes, because of the absence of nucleotide, and the use of Nb35 and scFV16 that stabilizes the conformation of the switch II loop and the αN-Gβγ interface respectively (*2, 9*). The recruitment of Gα_s_ to D1R was completely abolished, when N23, I26, E27 and L30 in αN were mutated to alanine to disrupt the αN and Gβγ interface **(Fig. 4G)**. Previous studies have shown that although αN truncations of Gα reduce the binding affinity between Gα and Gβγ, the truncated Gα could still interact with Gβγ (*37*). These data suggest that Gβγ contributes to the initial G protein coupling to the receptor partially by stabilizing the conformation of αN. Direct interactions between Gβγ and receptors that are observed in many structures of GPCR-G protein complexes are involved in G protein coupling as well (*8*). To further support our structural findings, we analyzed the effect of mutations that favor a GTP-bound conformational state on G_s_ dissociation kinetics using NanoBiT-based G protein dissociation assay. In the GDP-bound D1R complex, Y37 in αN makes a hydrogen bond with D240 in Gα, while in the GTP-bound D1R complex, the movement of αN disrupts this hydrogen bond **(Fig. 4H)**. As expected, Y37F mutation that disrupts its hydrogen bond with D240 and favors the GTP-bound state had little influence on Gs recruitment **(fig. S6A)** but led to a faster Gs dissociation rate catalyzed by D1R **(fig.S6B and Fig. 4, I and J)**. In conclusion, the conformational changes of the switch II region and αN serve as molecular basis for the GTP-dependent dissociation of Gβγ from Gα, and of G protein from receptors.

**Fig. 4.**
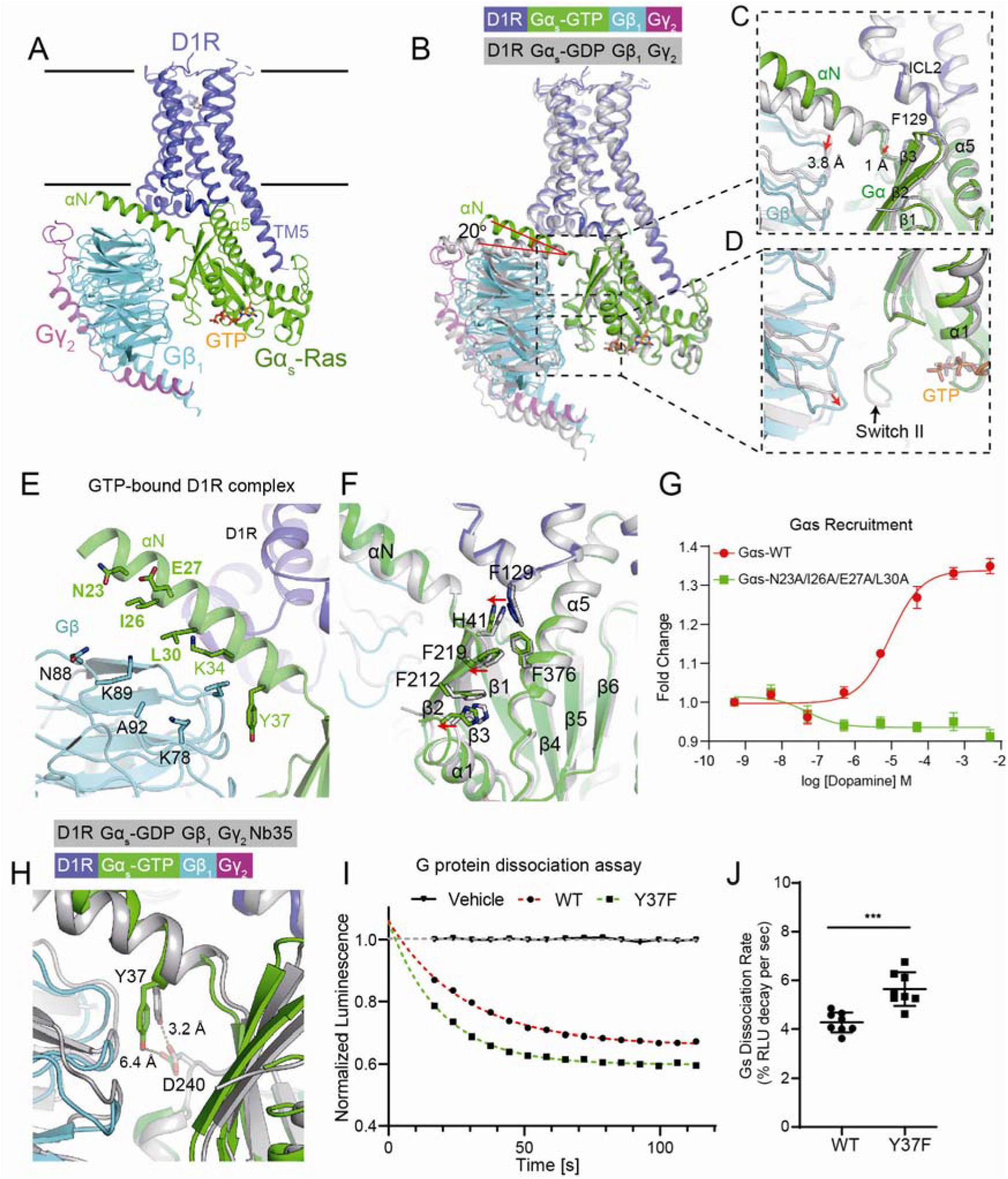
Structural changes of Gα upon receptor engagement after the exchange of GDP for GTP. **(A)** Overall structure of the GTP-bound D1R-mini-Gs complex with Nb35 dislodged. **(B)** Structural overlay of the GDP and GTP-bound D1R-miniGs complex without Nb35 bound. The αN is tilted 20° towards the receptor upon GTP binding. **(C and D)** Close-up view of conformational changes of the switch II, Gβγ and the αN-β1 hinge induced by GTP binding. **(E)** Interface of αN-Gβγ in the GTP-bound D1R-G protein complex**. (F)** Conformational differences between the GDP- and GTP-bound D1R complex. **(G)** Disruption of the αN-Gβγ interface abolishes G protein recruitment, as revealed by NanoBiT G protein recruitment assay using D1R-SmBiT and Gα_s_-LgBiT. **(H)** GTP binding disrupts the hydrogen bond between Y37 and D240 in Gα_s_. **(I)** G protein dissociation curve of Gα_s_ wild type and Y37F mutant at a saturated concentration of dopamine measured by NanoBiT dissociation assay. **(J)** Significance analysis of Gs dissociation rate of Gα_s_ wild type and Y37F mutant from eight independent experiments.

In summary, our data provide structural view of the entire GPCR-G protein coupling events, including initial G protein engagement by the receptor, receptor-mediated GDP release and GTP-dependent complex dissociation **(Fig. 5)**. The different binding poses of dopamine arising from variable GPCR-G protein interfaces among different studies provide further evidence of allosteric coupling from downstream effectors to ligand-binding pocket in GPCRs (*22*). We identified a prevalent sequence motif in TM5 of Gs-coupled receptors that plays an important role in determining G protein selectivity. The structure of the GDP-bound D1R-G protein complex reveals conformational steps of G protein activation by GPCR and critical regions for initial release of GDP. AHD domain is invisible in the most structures of GPCR-G protein complexes in the nucleotide-free state because of its high flexibility after the separation of AHD from Ras that occurs at the early stage of coupling events, even without receptor binding (*6*). Therefore, the conformational state of the GDP-bound complex captured here using mini-G protein that lack the AHD domain may represent an intermediate state of G protein upon receptor binding after AHD domain opening prior to GDP release but not the pre-coupled state where the α5 helix likely adopts a different configuration from our structures (*13*). Moreover, structural findings in the GTP-bound D1R complex highlight the important role of αN in G protein recruitment and GTP-dependent dissociation of G protein from the receptor. Taken together, our studies further advance our mechanistic understanding of G protein activation by GPCRs.

**Fig. 5.**
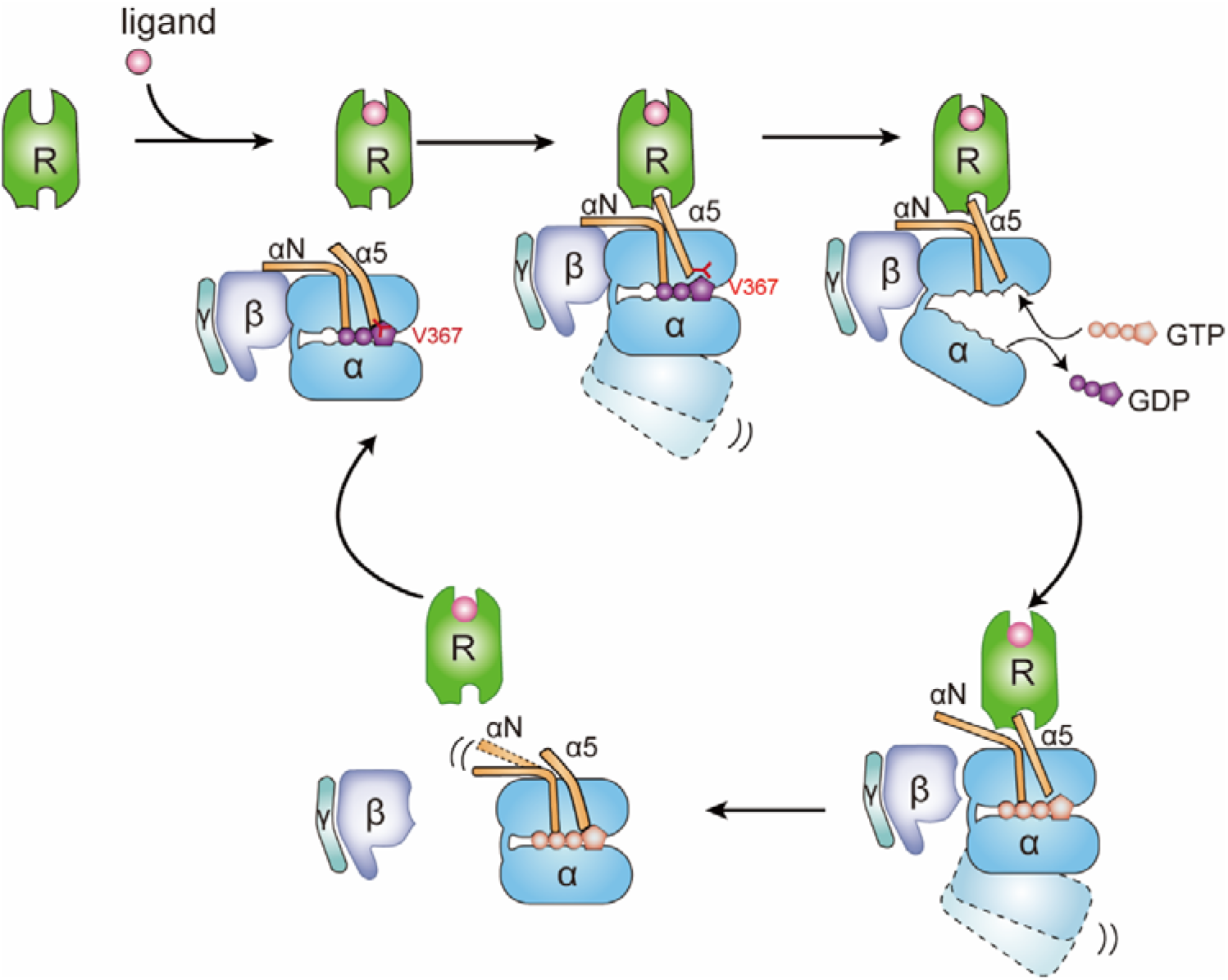
A model for the Gs activation by D1R. G protein engagement by the receptor causes the rotational and translational movement of α5, which leads to the upward movement of V367 and structural rearrangement of α1. These conformation changes altogether cause the separation of AHD and Ras domain and weaken the GDP binding affinity, leading to GDP release. Subsequent GTP binding results in the conformational change of αN and switch II, accounting for the dissociation of Gβγ from Gα.

## METHODS

### Cloning and expression of DR1-miniGs fusion protein

The human wild-type full-length D1R gene was cloned into a pcDNA3.1(+) vector (Thermo Fisher Scientific) with the signal peptide substituted by that of hemagglutinin (HA), and expressed with an N-terminal Flag tag and a C-terminal mini-Gα_s_399 fusion protein. 3C protease site was introduced between D1R and mini-Gα_s_ protein. Plasmids expressing fusion protein were transiently transfected into Expi293F cells (Thermo Fisher Scientific) using polyethyleneimine (Polysciences), when cells reached a density of 1.5 million per mL. 5 mM sodium butyrate and 3 mM valproic acid were added into the culture 18 h post-transfection, and cells were shaken for another 30 h before harvest by centrifugation at 1000 g for 10 min.

Cells were lysed in hypotonic buffer (25 mM HEPES-NaOH pH 7.6, 50 mM NaCl and 100 μM dopamine) using glass dounce tissue grinder. Membrane was pelleted by centrifugation at 60000g at 4 °C for 1 h and homogenized in solubilization buffer containing 25 mM HEPES pH 7.6, 150 mM NaCl, 0.5% LMNG (Anatrace), 0.1% cholesteryl hemisuccinate (CHS, Anatrace), 10 μM dopamine (Sigma-Aldrich) using dounce. Sample was mixed for 2h at 4 °C. After centrifugation to remove the debris, the supernatant supplemented with 2 mM CaCl_2_ was loaded onto anti-Flag antibody affinity resin by gravity flow. The resin was washed extensively with at least 10 column volume of wash buffer containing 25 mM HEPES, pH 7.6, 150 mM NaCl, 0.01% LMNG, 0.002% CHS, 2 mM CaCl_2_, 10□mM MgCl_2_, 2 mM KCl and 2□mM adenosine triphosphate, 10 μM dopamine. The receptor was eluted in elution buffer (25mM HEPES, 150 mM NaCl, 0.01% LMNG, 0.002% CHS, 5 mM EDTA, 0.1 mg/ml Flag peptide, 10 μM dopamine). The protein sample was concentrated by ultrafiltration and incubated with PNGaseF (New England Biolabs) overnight.

### Complex assembly

His6-tagged human Gβ_1_ and Gγ_2_ with C68S mutation was expressed in insect cell using the Bac-to-Bac Baculovirus expression system (Invitrogen) and purified as previously described (*38*). Nb35 was expressed in *Escherichia coli strain* BL21 (DE3) and purified as previously reported (*2*). For the D1R-miniGα_s_-Gβ_1_γ_2_-Nb35 complex assembly and purification, purified D1R-mini-Gα_s_ fusion protein, Gβ_1_γ_2_ and Nb35 were mixed in a 1:1.2:1.2 molar ratio and added with 2 mM MgCl_2_ and apyrase. Nb35 was not included for the D1R-mini-Gs-Gβ_1_γ_2_ complex assembly. After incubation at 4 °C overnight, the protein complex was further purified with superose 6 10/300 to remove the excess Gβ_1_γ_2_ and Nb35 in buffer containing 25 mM HEPES pH 7.6, 150 mM NaCl, 0.01% LMNG, 0.002% CHS, and 10 μM dopamine. The complex peak were pooled and concentrated to 4 mg/ml for cryo-EM analysis.

### Cryo-EM sample preparation and data collection

3.0 μl of purified complex was applied to glow-charged 300 mesh holey carbon grid (Quantifoil Au R1.2/1.3). Grids were blotted for 3.0-4.0 s at a blotting force of 4 and vitrified using a Vitrobot MarkIV (Thermo fisher Scientific) with chamber maintained at 8 °C and 100% humidity. For the nucleotide-bound complex, 1 mM GDP or GTP and 2 mM MgCl_2_ were added to the protein sample prior to grid preparation using the same condition as above. Cryo-EM movies were collected on a Titan Krios (Thermo Fisher Scientifc) operated at 300 kV and equipped with a BioQuantum GIF/K3 direct electron detector (Gatan) in a superresolution mode at a nominal magnification of ×64,000. Each movie stack was collected as 32 frames with a total dose of 50 e^-^/Å^2^ for 2.56 s. Cryo-EM data collection parameters for all protein samples are summarized in Table S1.

### Data processing

For the nucleotide-free D1R-mini-Ga_s_-Gβ_1_γ_2_-Nb35 complex, a total of 2320 movie stacks were collected and subjected to motion correction with 2× binned to a pixel size of 1.087 Å using MotionCor2(*39*). Contrast transfer function (CTF) estimation was performed using patch-based CTF estimation in cryoSPARC (*40*). 3,876,379 particles were auto-picked using the Blob picker in cryoSPARC. These particles were split into three groups extracted in a 180-pixel box and subjected to 2D classification in cryoSPARC. Particles with good 2D class average were combined and run through the next round of 2D classification. *Ab-initio* reconstruction with five classes using 1,045,088 particles was performed in cryoSPARC and subjected to heterogeneous refinement. Particles from classes with clear secondary structure were selected and run through another round of *Ab-initio* reconstruction with six classes and subsequent heterogeneous refinement. Two classes with high resolution and clear transmembrane helices were combined and applied to non-uniform refinement in cryoSPARC, resulting in a map with global resolution of 3.1 Å.

For the GDP-bound D1R-mini-Ga_s_-Gβ_1_γ_2_-Nb35 complex, a total of 601 movies were collected, and similar procedure was performed as above. In brief, *ab-initio* reconstructions with five classes using 317,029 particles yield two good classes with clear secondary structure, accounting for 65.3% of total particles. The two classes were combined and subjected to non-uniform refinement, yielding a map with global resolution of 3.1 Å.

For the GDP-bound D1R-mini-Gα_s_-Gβ_1_γ_2_ complex, 448,009 particles with good 2D class average from 681 movies were extracted in a 180-pixel box in cryoSPARC and exported into RELION format using csparc2star.py script from UCSF pyem package (*41*). These particles were used for 3D classification in RELION (*42*). One class accounting for 46.3% particles showing a well-defined structure was selected and imported back to cryoSPARC and run through non-uniform refinement to yield a map at 3.5 Å resolution.

For the GTP-bound D1R-mini-Gα_s_-Gβ_1_γ_2_-Nb35 complex, Particles from 1242 movies were subjected to two round of 2D classification by cryoSPACR and one round of 2D classification by RELION, yielding 628,083 good particles. 3D classification was performed in RELION, resulting in one good class accounting for 49.5% particles. The next round of 3D classification yielded two classes with clear transmembrane helices, one with Nb35 occupied and one with Nb35 dislodged. For the complex without Nb35, we performed 3D refinement with mask excluding micelle. For the complex with Nb35, particles were imported to cryoSPARC and run through non-uniform refinement to yield a map at 3.6 Å resolution. Resolutions are reported based on the gold standard Fourier shell correlation (FSC) at the 0.143 criterion.

All cryo-EM maps were post-processed by DeepEMhancer to improve their interpretability (*43*).

### Model building

A homology model of D1R was generated using SWISS-MODEL server (*44*) with activated structure of β1AR (PDBID: 7JJO) as a template and was docked into the EM density map along with miniGs-Nb35 structure in Chimera (*45*). The model was manually built in COOT (*46*) and refined with *Phenix* (*47*). Initial restraints for dopamine, GDP and GTP were generated using eLBOW in *phenix*. If the side chain density is too poor to assign a conformation, we temporarily chop the side chain while keeping sequence information. Model was validated using Molprobity (*48*) and EMRinger (*49*). Model-to-map FSC curves were calculated in Phenix. Structure figures are prepared with Pymol and Chimera. Detailed structure statistics are summarized in Table S1.

### cAMP accumulation assay

The human full-length D1R gene was cloned into pcDNA3.1(+) vector with an N-terminal flag tag. All point mutations are introduced by the QuikChange method. HEK293 cells stably expressing the GloSensor biosensor were plated into six-well plate in Dulbecco’s modified Eagle’s medium (DMEM, Gibco) supplemented with 10% fetal bovine serum (FBS, Gibco), penicillin and streptomycin, and transfected with wild-type or mutated D1R plasmids using polyethylenimine. After transfection, cells were incubated at 37 °C with 5% CO_2_ for 24 h. Then cells were collected and seeded in a tissue culture-treated, white, and clear-bottom 96-well plate. After incubation for another 24 h, culture medium were removed, and equilibration medium (CO_2_-independent medium, 10% FBS and 1% D-luciferin) were added to each well. Cells were incubated at room temperature for 2 h before treatment with increasing concentration of dopamine. The luminescence signal was measured in 10 min after the addition of dopamine and plotted as a function of dopamine concentration using nonlinear regression with GraphPad Prism 8 (GraphPad Software). EC50 indicates the concentration of ligand which can produce 50% of the maximum luminescence signal. Each measurement was repeated in three independent experiments, each in triplicate. Significance was calculated by two-tailed student’s t-test.

### NanoBiT Gs dissociation assay

NanoBiT-based Gs dissociation assay was performed as previously described (*50*). The large fragment (LgBiT) and small fragment (SmBiT) that comprise a catalytically active luciferase were fused to the AHD domain of Gα_s_ (Gα_s_-LgBiT) and the N-terminus of Gγ_2_ with a C68S mutation (SmBiT-Gγ2), respectively. HEK293T cells were seeded in a six-well plate using the same DMEM medium as above. 200 ng D1R, 100 ng Gα_s_-LgBiT, 500ng Gβ_1_, 500 ng SmBiT-Gγ_2_ and 100ng RIC8B were transfected into cells using polyethylenimine solution, when cells reach 80% confluency. After 1 day incubation, cells were washed with Dulbecco’s PBS and suspended in 3 ml HBSS reaction buffer (HBSS supplemented with 0.01% BSA and 5 mM HEPES, pH 7.4). Coelenterazine was added to cell suspensions at a final concentration of 10 μM. Cells were seeded into 96-well plate with 1 × 10^5^ cells per well in 95 μl of HBSS reaction buffer. After incubation at room temperature for 1 h, baseline luminescence signals were measured using luminescent microplate reader (Tecan, Spark). 5 μl of increasing concentration of dopamine (20× of final concentrations) diluted in HBSS reaction buffer was added to cells. Luminescence signals were measured in 3-5 min after ligand addition and normalized over baseline signal. The resulting fold-changes are plotted as a function of concentrations of dopamine using a three-parameter sigmoidal concentration-response model built in Prism 8.0.

To calculation the dissociation speed at a concentration of dopamine producing saturated luminescence, the plate was immediately read at an interval of 6.8 s with an accumulation time of 0.5 s per read for 2 min following ligand addition. The luminescence signal was normalized to the baseline count. The normalized signal was fitted using one-phase dissociation model built in Prism 8.0. The dissociation speed K represented decreased luminescence per second.

### NanoBiT G protein recruitment assay

For monitoring recruitment of Gβ_1_γ_2_, LgBiT and SmBiT were fused with the C-terminus of D1R and the N-terminus of Gβ_1_ to yield D1R-LgBiT and SmBiT-Gβ_1_ fusion proteins, respectively. Plasmid mixtures containing 200 ng D1R-LgBiT, 100 ng Gα_s_, 500 ng SmBiT-Gβ_1_, 500 ng Gγ_2_C68S and 100 ng RIC8B were transfected into HEK293T cells.

For directly monitoring recruitment of Gα, D1R-SmBiT containing D1R fused to SmBiT at its C-terminus, Gα_s_-LgBiT, Gβ_1_ and Gγ_2_C68S were expressed with RIC8B in HEK293T cells using same amount of plasmids as above.

For mini-Gs recruitment assay, LgBiT-mini-Gα_s_ consisting of mini-Gα_s_399 (*11*) fused to LgBiT at its N-terminus and D1R-SmBiT were coexpressed in HEK293T cells.

Similar procedures were performed as G protein dissociation assay. In brief, luminescence signals were measured in 3-5 min following addition of increasing concentration of dopamine, and normalized to baseline signal. The resulting fold changes were fitted by non-linear regression using Prism.

### GTP turnover assay

Human Gα_s_ and its mutants used for the assay were expressed and purified from bacteria. Gα_s_ (residue 7-394) was cloned into pET28a vector with an N-terminal His_6_-SUMO-Flag tag. All point mutations in Gα were introduced using Quikchange method. The plasmids were transformed into *Escherichia coli* BL21 (DE3). The transformed bacteria were cultured in LB medium supplemented with 50 μg/ml kanamycin at 37 °C to an OD_600_ value of 0.8, and were shaked at 25 °C overnight following addition of 500 μM β-D-thiogalactopyranoside (IPTG). After harvest by centrifugation, cells were resuspended in lysis buffer (20mM HEPES pH 7.4, 300 mM NaCl, 2 mM MgCl_2_, 10 μM GDP, 100 μM TCEP, 15% glycerol) and lysed by sonication. Cell lysate was supplemented with ULP1 to cleave His_6_-SUMO tag, and flag-tagged Gα_s_ was purified by M1 Flag affinity chromatography. Resin was washed with wash buffer containing 20mM HEPES pH 7.4, 100 mM NaCl, 2 mM MgCl_2_, 2 mM CaCl_2_, 10 μM GDP, 100 μM TCEP and proteins were eluted with elution buffer containing 20mM HEPES pH 7.4, 100 mM NaCl, 2 mM MgCl_2_, 10 μM GDP, 100 μM TCEP, 5 mM EDTA, 0.1 mg/ml Flag peptide. The eluted Gα_s_ was incubated with 1.2-fold molar excess of Gβ_1_γ_2_ at 4 °C for 1 hour. The assembled complex was further purified by size exclusion chromatography on a Superdex 200 10/300 Increase column in buffer containing 20 mM HEPES pH 7.4, 100 mM NaCl, 5 mM MgCl_2_, 0.03% DDM, 10 μM GDP, 100 μM TCEP. Peak fractions were pooled and concentrated to 1 mg/ml for GTP turnover assay.

The GTP turnover assay was performed as previously described (*31*). 1 μM DDM-solubilized D1R was incubated with 200 μM dopamine in buffer containing 20mM HEPES pH 7.4, 100 mM NaCl, 0.03% DDM for 60 min at room temperature. A final concentration of 10 μM GTP was added into D1R before mixing D1R with 500 nM G protein in buffer containing 20 mM HEPES, 100 mM NaCl, 20 mM MgCl_2_, 0.03% DDM, 200 μM TCEP and 1 μM GDP. After incubation for an indicated time, reconstituted GTPase-Glo reagent made according to the manufacture’s protocol (Promega) was added to the reaction and incubated for 30 min at room temperature. Luminescence was measured in 5 min following the addition of detection reagent at room temperature using Tecan Spark. The data was normalized to the initial count of Gs without addition of receptor and then analyzed using Prism 8. Significance was obtained by two-tailed student’s t-test with Welch’s correction.

## Acknowledgements

We thank Andrew C. Kruse for critical reading of this manuscript. We thank staff at Shuimu BioSciences for their help with cryo-EM data collection. All EM images were collected at Shuimu BioSciences.

## Funding

This work was supported by Chinese Ministry of Science and Technology, Beijing Municipal Science & Technology Commission (Z201100005320012) and Tsinghua University.

## Author contributions

X.T. purified and assembled the protein complex. X.T. and S.Z. collected cryo-EM data and performed cryo-EM data processing and model building. X.T. and S.C. did the cAMP accumulation assay. X.W. purified Gβγ protein. S.Z. wrote the manuscripts with input from all other authors.

## Competing interests

The authors declare no competing interests.

## Data availability

The atomic structures have been deposited at the Protein Data Bank (PDB) under the accession codes 7F0T, 7F1O, 7F1Z, 7F23, and 7F24. The EM maps have been deposited at the Electron Microscopy Data Bank (EMDB) under the accession numbers EMD-31404, EMD-31421, EMD-31425, EMD-31426 and EMD-31427.

## Supplementary Materials for

**Fig. S1.**
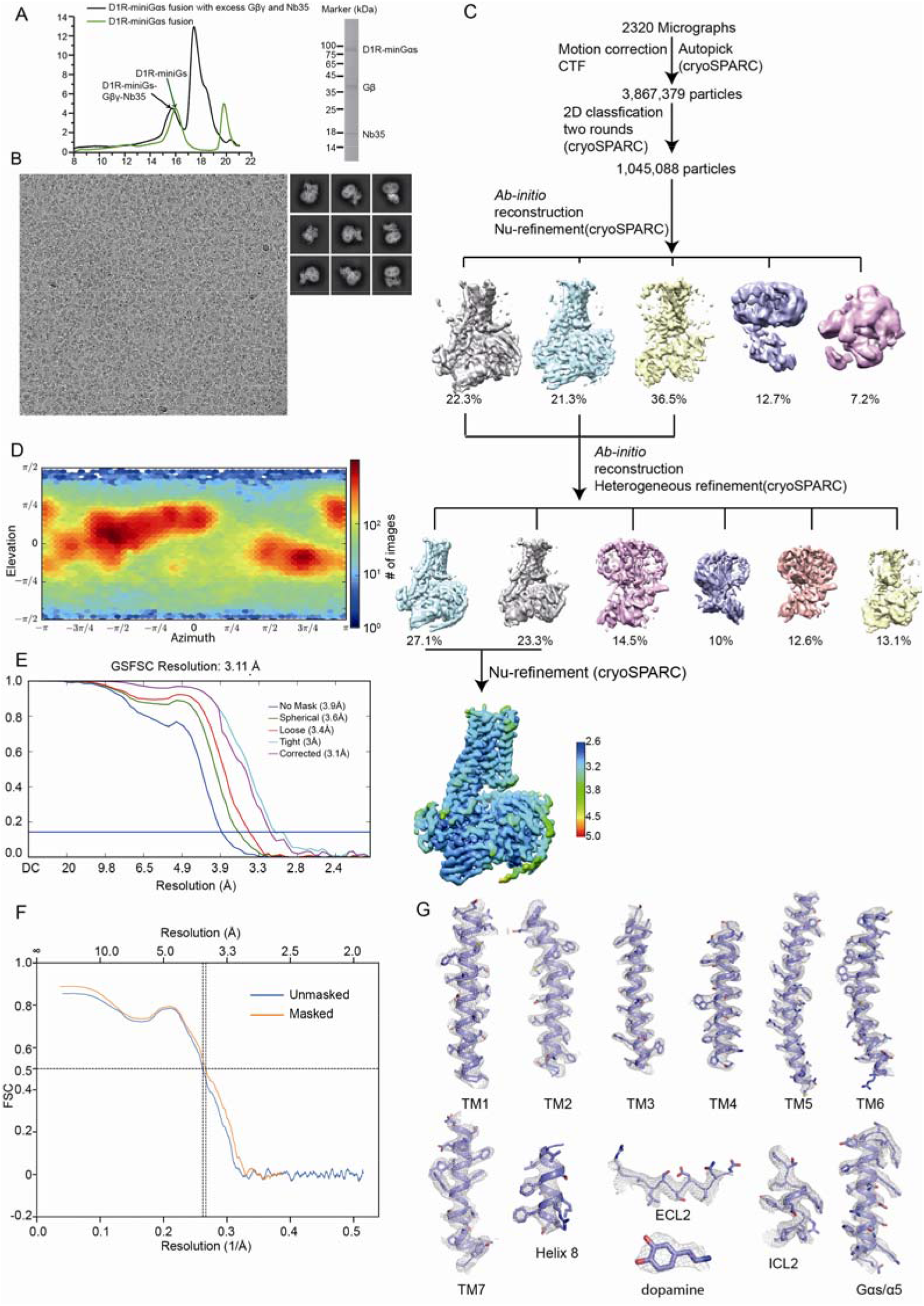
Cryo-EM data processing for the dopamine-bound D1R-mini-G_s_-Nb35 complex in the nucleotide-free state. **(A)** Size exclusion profiles of the D1R-miniGαs fusion protein and D1R-mini-Gα_s_-Gβγ-Nb35 complex (left), and SDS-PAGE of the D1R-mini-Gα_s_-Gβγ-Nb35 complex (right). **(B)** Representative cryo-EM micrograph (left) and 2D class average. **(C)** Cryo-EM workflow chart of data processing. **(D)** Angular distribution plot. **(E)** Gold standard FSC curves. (**F)** FSC of Model-to-map. **(G)** Representative EM density map of the D1R-mini-G_s_-Nb35 complex.

**Fig. S2.**
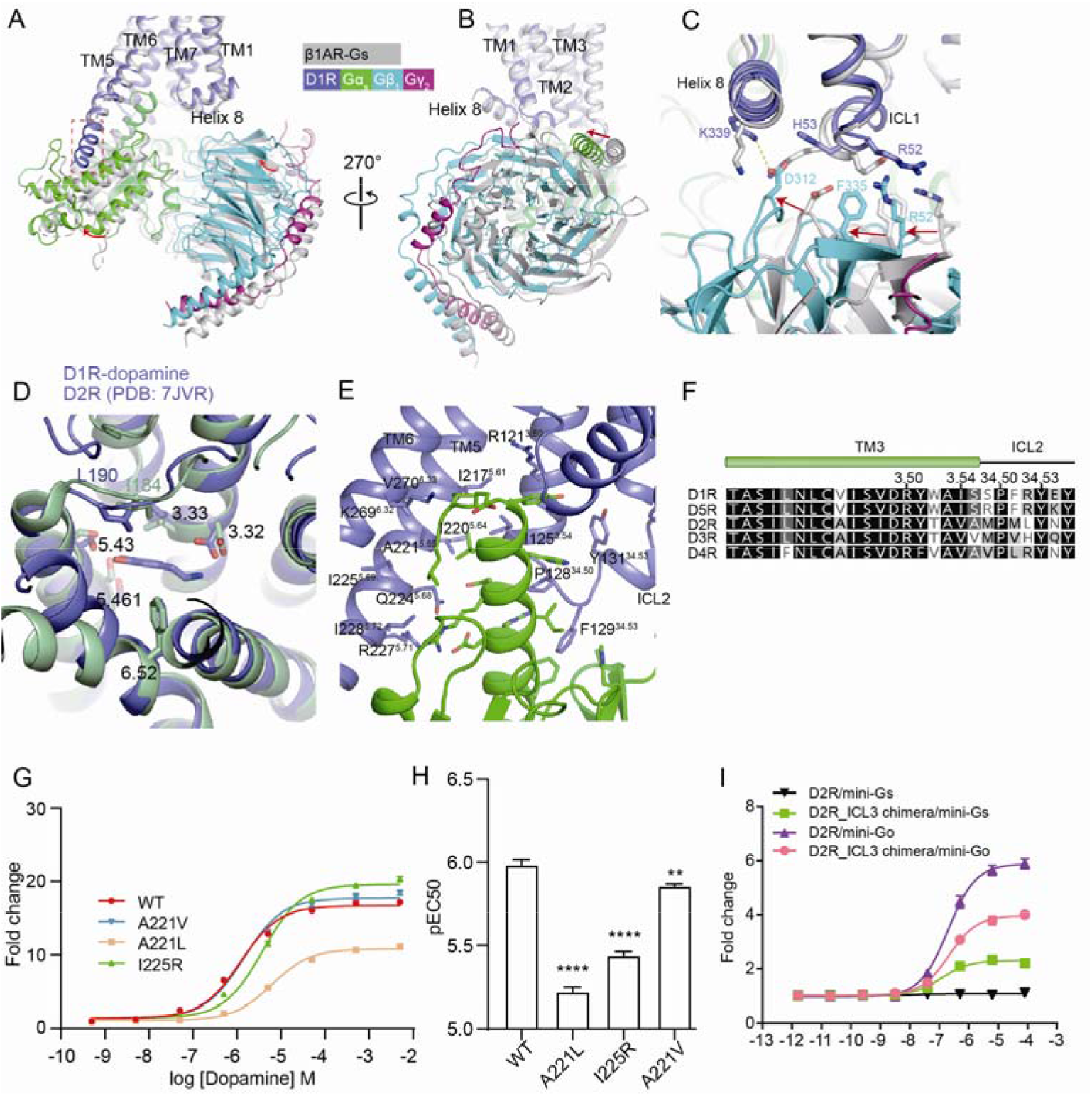
Structural analysis on the D1R-minGs-Nb35 complex in the nucleotide-free state. **(A and B)** Comparison of the structures of β1AR-G_s_ (PDB: 7JJO) and D1R-G_s_ complex without nucleotide bound in two orthogonal views. The extended TM5 in D1R-minG structure is boxed. **(C)** The detailed view of the receptor and Gβγ interface from D1R-G_s_ and β1AR-G_s_ complex. **(D)** Comparison of the dopamine binding pocket of D1R and D2R. Residues involved in binding dopamine were shown as sticks. **(E)** Interaction between D1R (blue) and the α5 of Gα (green). **(F)** Sequence alignment of TM3 and ICL2 from dopamine receptors. Residues involved in receptor binding were indicated by residue number above the alignment. **(G)** Effects of A221 and I225 mutations in D1R on G protein recruitment as evaluated by NanoBiT mini-G_s_ recruitment assay using D1R-SmBiT and LgBiT-mini-Gα_s_. **(H)** EC50 obtained from NanoBiT mini-G_s_ recruitment assay. Data indicate mean ± SEM from three independent experiments performed in triplicate. **(I)** NanoBiT mini-G_s_ recruitment results show that the ability of D2R to recruit Gα_s_ is significantly enhanced, when the ICL3 of D2R is replaced by that of D1R.

**Fig. S3.**
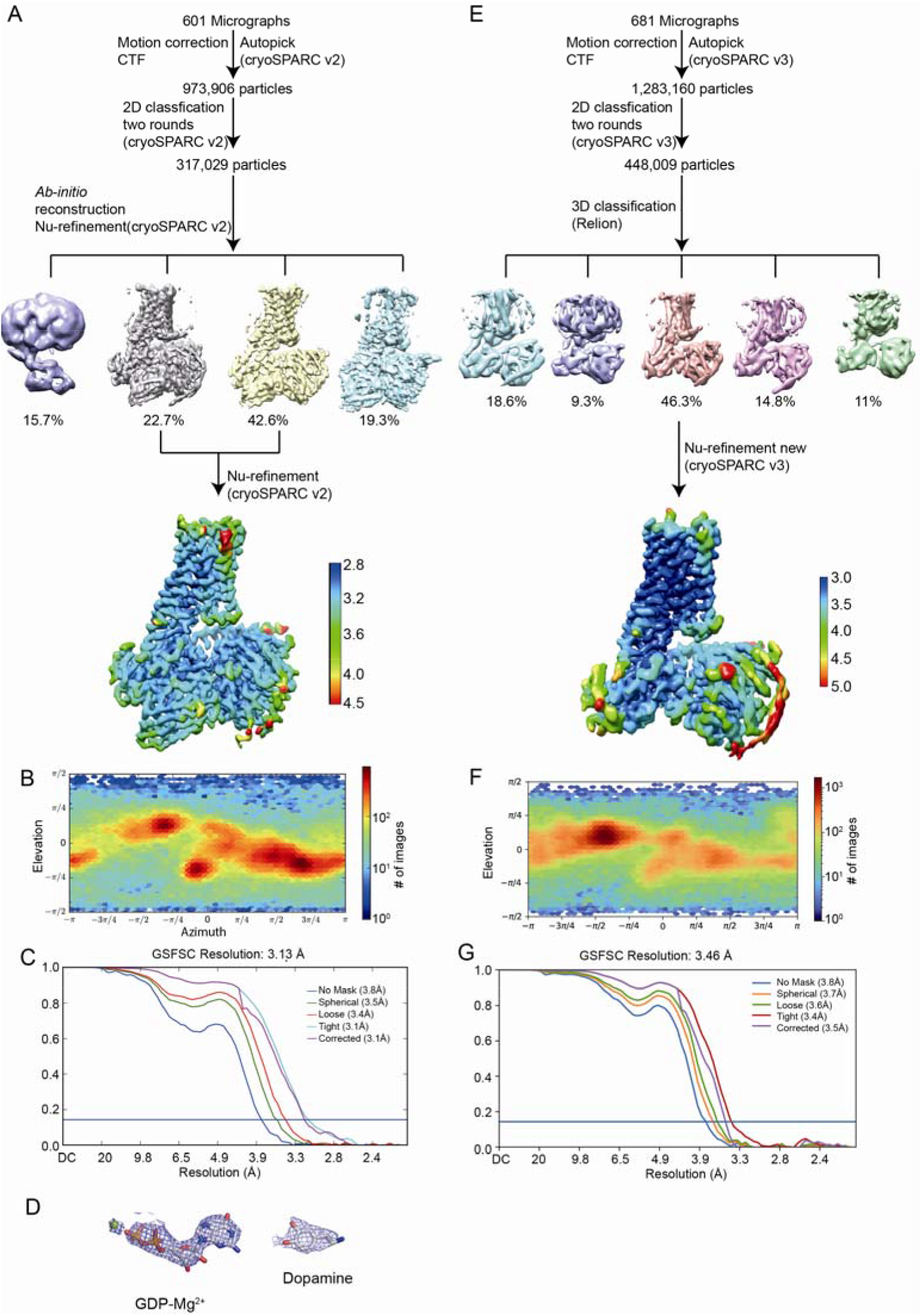
Cryo-EM data processing for the dopamine-bound D1R-mini-G_s_ complex in the presence of GDP. **(A)** Cryo-EM workflow chart for D1R-mini-G_s_-Nb35 complex with GDP-bound. **(B)** Angular distribution plot for D1R-mini-G_s_-Nb35 complex with GDP-bound. **(C)** Gold-standard FSC curve of D1R-mini-G_s_-Nb35 complex with GDP-bound. **(D)** EM density map of GDP and dopamine from the D1R-mini-G_s_-Nb35 complex with GDP-bound. **(E)** Cryo-EM workflow chart of the GDP-bound D1R-mini-G_s_ complex without Nb35. **(F)** Angular distribution plot for the GDP-bound D1R-mini-G_s_ complex without Nb35. **(G)** Gold-standard FSC curve of the GDP-bound D1R-mini-G_s_ complex without Nb35.

**Fig. S4.**
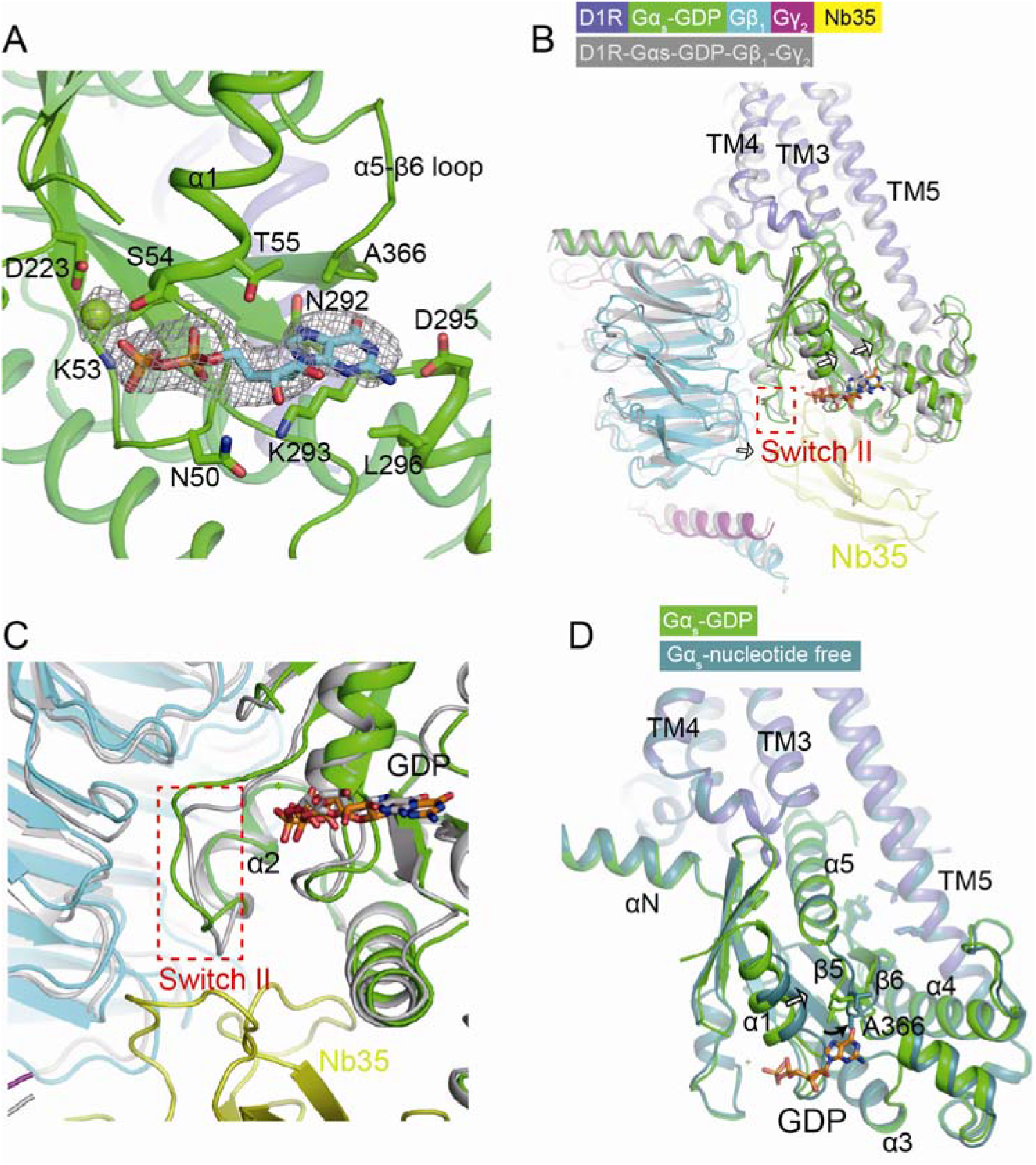
Structural analysis of the GDP-bound D1R-mini-G_s_ protein complex. **(A)** View of GDP and Mg^2+^ in the structure of the GDP-bound D1R-miniGs-Nb35 complex. **(B)** Comparison of structures of GDP-bound D1R-mini-G_s_ with and without Nb35 binding. Receptors were aligned. **(C)** The effect of Nb35 on the conformational change of switch II in the D1R-mini-G_s_ complex. **(D)** Comparison of structures of D1R-mini-G_s_ in the nucleotide-free state and the GDP-bound state.

**Fig. S5.**
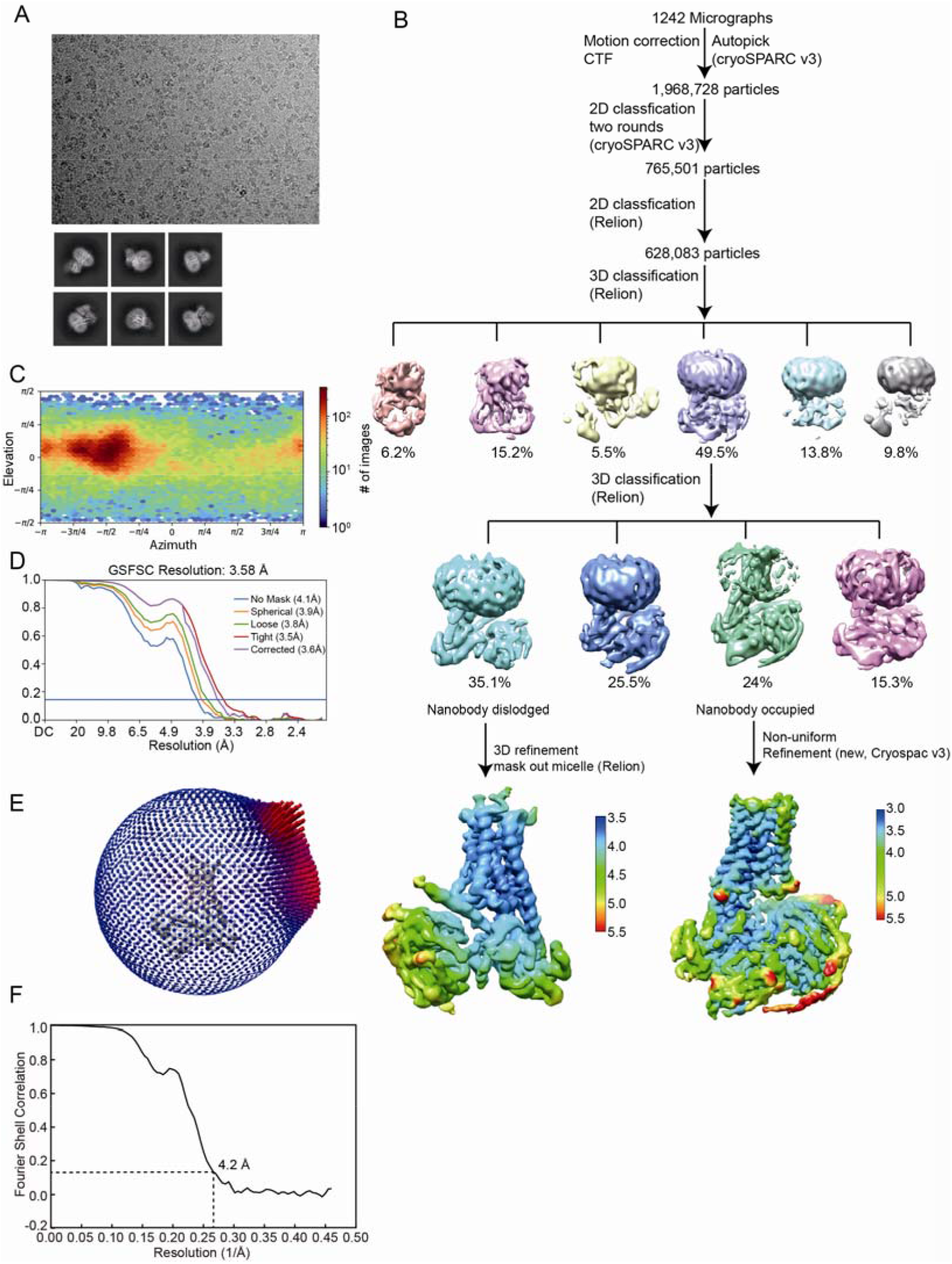
Cryo-EM workflow chart for the D1R-mini-G_s_-Nb35 complex with GTP-bound**. (A)** Representative micrograph (up) and 2D class average (bottom) for the D1R-mini-G_s_-Nb35 complex with GTP-bound. **(B)** Cryo-EM workflow chart of the GTP-bound D1R-mini-G_s_-Nb35 complex**. (C and D)** Angular distribution and FSC curve of the GTP-bound D1R-mini-G_s_-Nb35 complex with Nb35 occupied. **(E and F)** Angular distribution and FSC curve of the GTP-bound D1R-mini-G_s_-Nb35 complex with Nb35 dislodged.

**Fig. S6.**
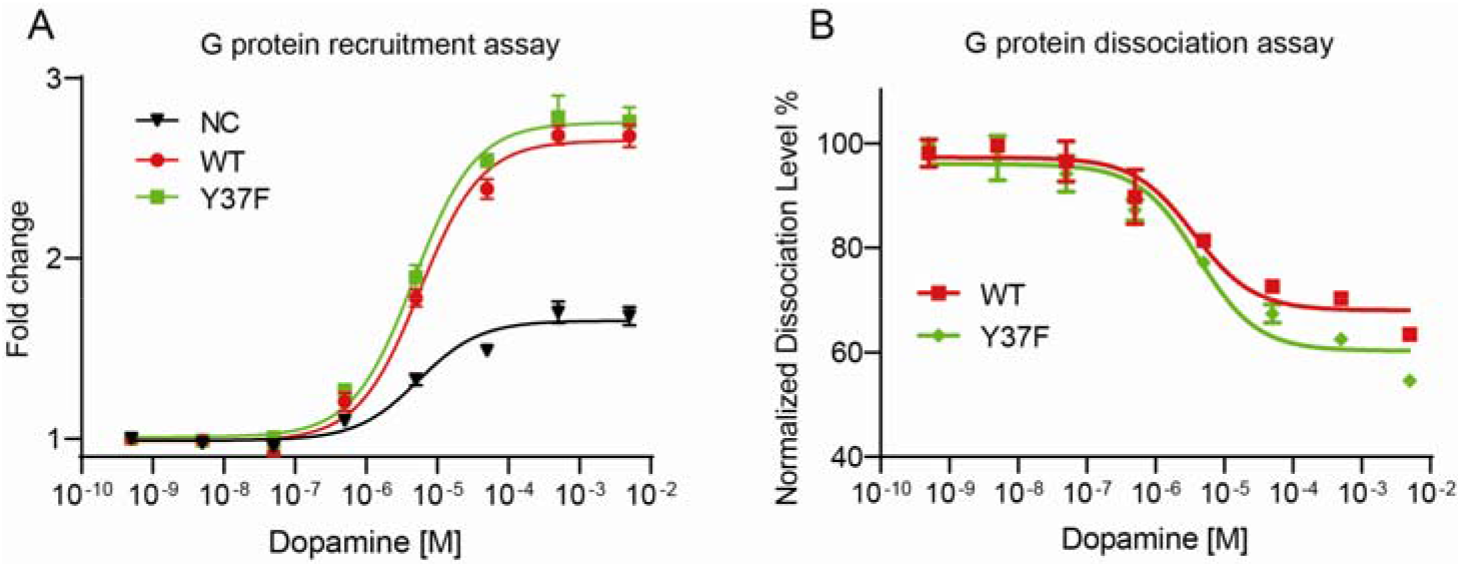
The effect of Gαs-Y37F mutation on G protein recruitment and dissociation. (A) NanoBiT G protein recruitment assay using D1R-LgBiT and SmBiT-Gβ_1_. (B) NanoBiT G protein dissociation assay.

**Table S1.**
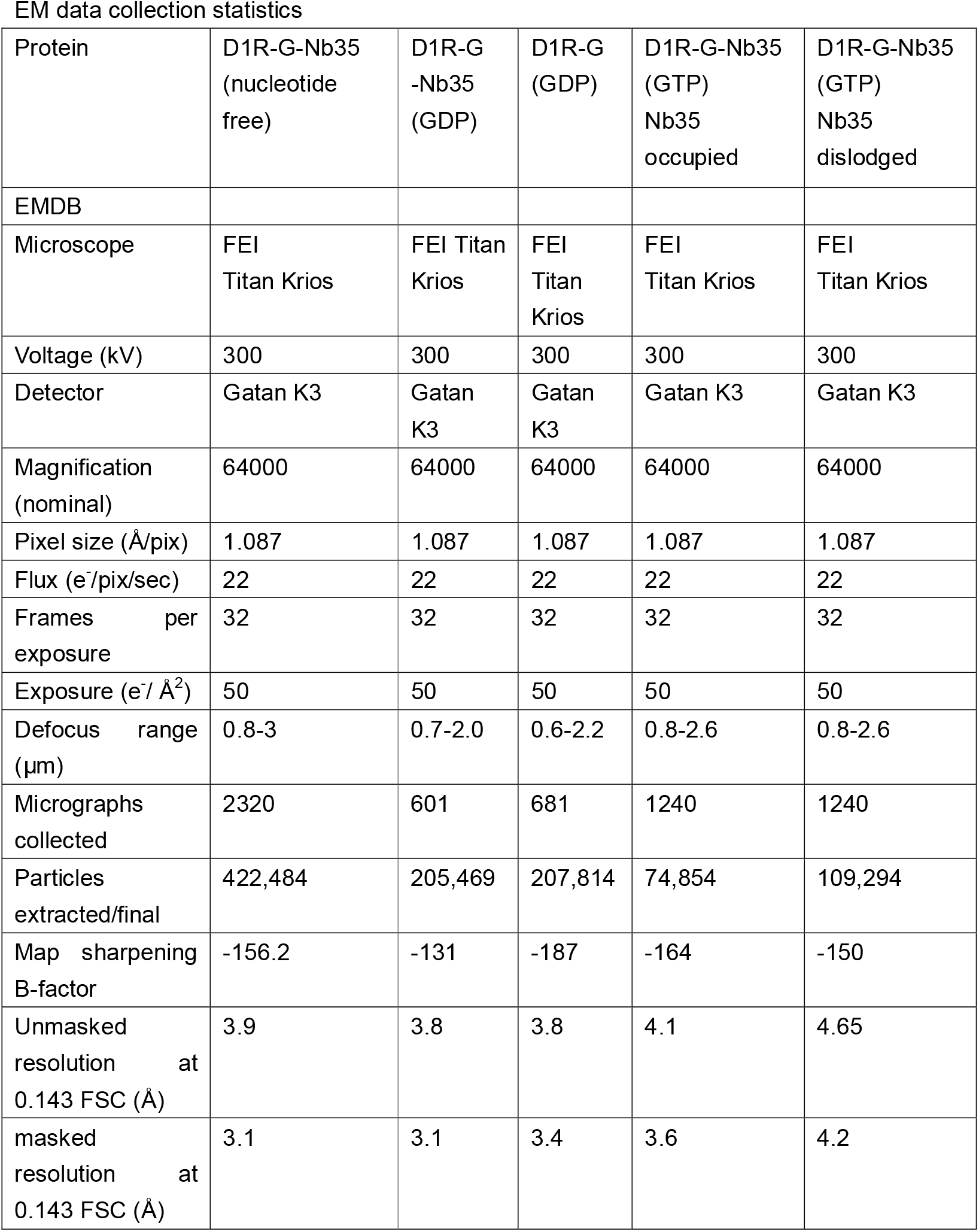
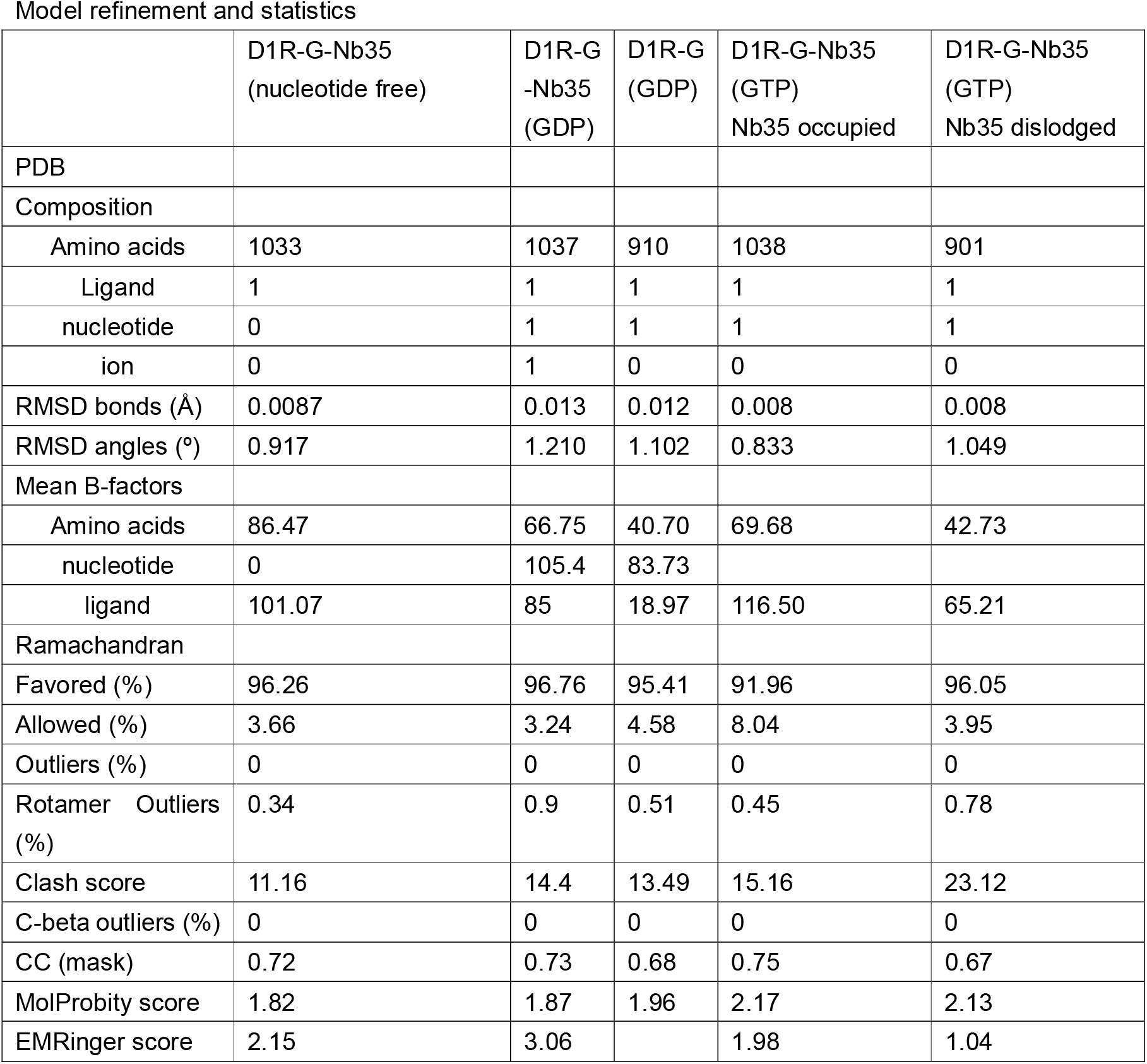
Cryo-EM data collection and refinement statistics.

## Notes

### Competing Interest Statement

The authors have declared no competing interest.

